# Next-generation sequencing in neuropathological diagnosis of infections of the nervous system

**DOI:** 10.1101/039222

**Authors:** Steven L. Salzberg, Florian P. Breitwieser, Anupama Kumar, Haiping Hao, Peter Burger, Fausto J. Rodriguez, Michael Lim, Alfredo Quiñones-Hinojosa, Gary L. Gallia, Jeffrey A. Tornheim, Michael T. Melia, Cynthia L. Sears, Carlos A. Pardo

## Abstract

**Objective:** To determine the feasibility of next-generation sequencing (NGS) microbiome approaches in the diagnosis of infectious disorders in brain or spinal cord biopsies in patients with suspected central nervous system (CNS) infections.

**Methods:** In a prospective-pilot study, we applied NGS in combination with a new computational analysis pipeline to detect the presence of pathogenic microbes in brain or spinal cord biopsies from ten patients with neurological problems indicating possible infection but for whom conventional clinical and microbiology studies yielded negative or inconclusive results.

**Results:** Direct DNA and RNA sequencing of brain tissue biopsies generated 8.3 million to 29.1 million sequence reads per sample, which successfully identified with high confidence the infectious agent in three patients, identified possible pathogens in two more, and helped to understand neuropathological processes in three others, demonstrating the power of large-scale unbiased sequencing as a novel diagnostic tool. Validation techniques confirmed the pathogens identified by NGS in each of the three positive cases. Clinical outcomes were consistent with the findings yielded by NGS on the presence or absence of an infectious pathogenic process in eight of ten cases, and were non-contributory in the remaining two.

**Conclusions:** NGS-guided metagenomic studies of brain, spinal cord or meningeal biopsies offer the possibility for dramatic improvements in our ability to detect (or rule out) a wide range of CNS pathogens, with potential benefits in speed, sensitivity, and cost. NGS-based microbiome approaches present a major new opportunity to investigate the potential role of infectious pathogens in the pathogenesis of neuroinflammatory disorders.

## Introduction

Ascertainment of the etiology of inflammatory disorders of the central nervous system (CNS) represents a major challenge in the clinical setting, with recent studies estimating that more than 50% of cases go undiagnosed^1^. Next generation sequencing (NGS) and metagenomic technologies present a major new opportunity to investigate the potential role of infection in the pathogenesis of neuroinflammatory disorders. This technology can provide a view of the transcriptome of the host tissue as well as capture microbial genomes (i.e., bacteria, fungi, and viruses) that reside in the tissue niche^2^–^4^. A major challenge to this new approach is the presence of non-pathogenic microbes, which are present on and beneath the skin and in multiple organs of the human body. Until recently, most sequence-based pathogen identification studies have focused on targeted capture of the 16S rRNA gene that is exclusive to prokaryotes. The availability of very high throughput, low-cost sequencing now makes it possible to sequence tissue samples deeply enough to detect pathogens in a background dominated by human DNA. Deep sequencing of total DNA or RNA provides an unbiased approach that can detect even rare components of the microbiome^5^. This strategy has recently been used to diagnose cases of encephalitis^6^, neuroleptospirosis^7^, astrovirus^8–10^, and bornavirus^11^ infection, but other than these isolated cases, the utility of NGS for clinical or pathological diagnosis has yet to be established.

We report here a pilot-prospective study of the use of unbiased NGS to assist in the diagnosis of neuroinflammatory disorders suspected to be associated with infections of the

CNS. In a series of ten patients in whom brain or spinal cord biopsies were taken as part of standard care, and in whom an infectious cause was suspected, we applied NGS in combination with a new computational analysis pipeline to detect the presence of pathogenic microbes. The computational pipeline includes a very rapid method for identifying pathogen species based on short, next-generation sequencing (NGS) reads, as short as 100 base pairs (bp) in length^12^.

## Methods

### Patients, biopsy handling and sequencing

CNS tissues were obtained prospectively from biopsies performed during diagnostic assessment of 10 patients with neuroinflammatory disorders suspected to be associated with infection (Table 1). All biopsies but one were obtained at Johns Hopkins Hospital and one biopsy was referred for consultation to the Division of Surgical Neuropathology. Tissue was collected under a research protocol approved by the Johns Hopkins University School of Medicine IRB. Fresh frozen biopsy tissues from eight cases were sequenced immediately after biopsy over the course of the study. Other two biopsy samples were from paraffin-processed tissues (See Supplemental Methods).

**Table 1.**
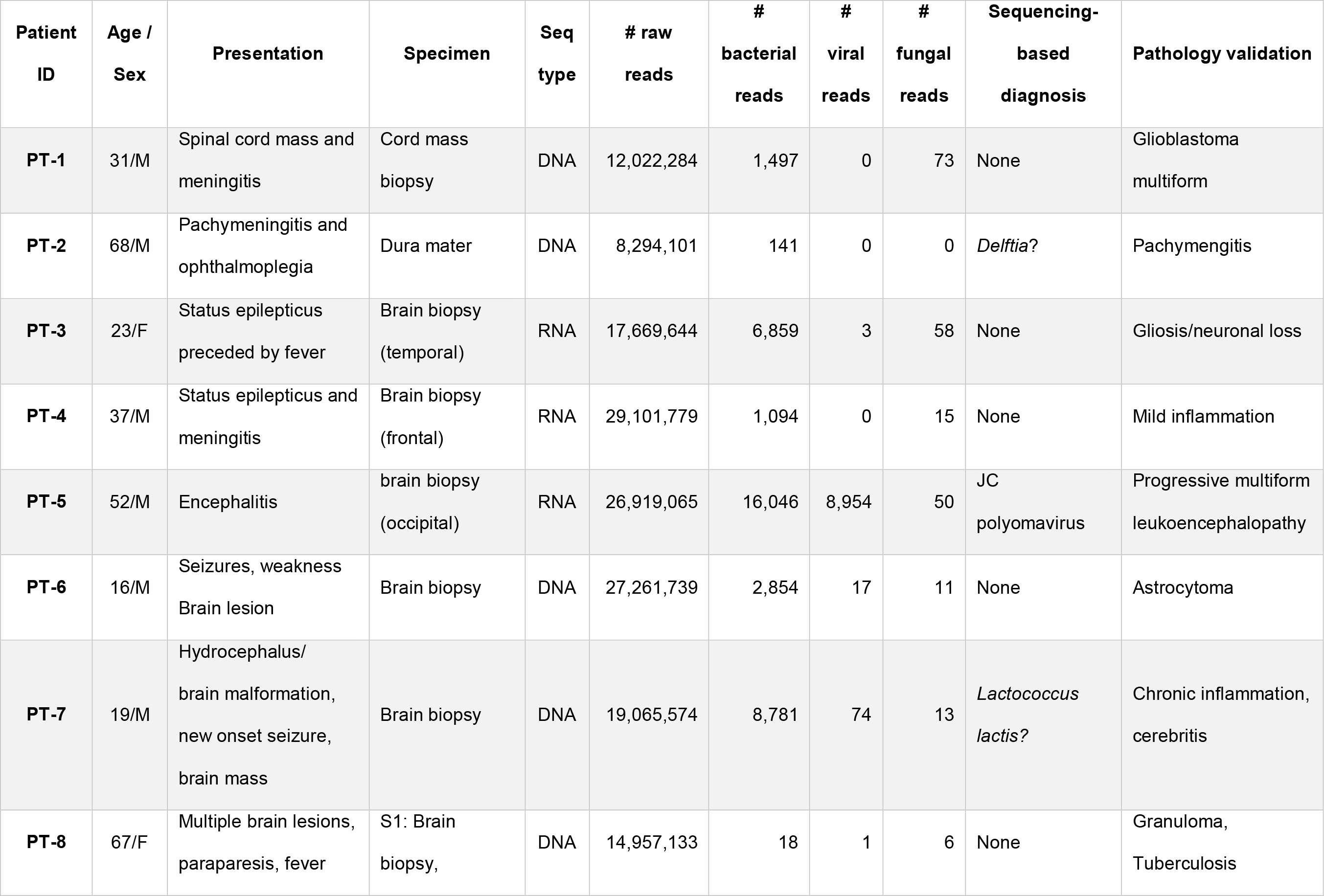
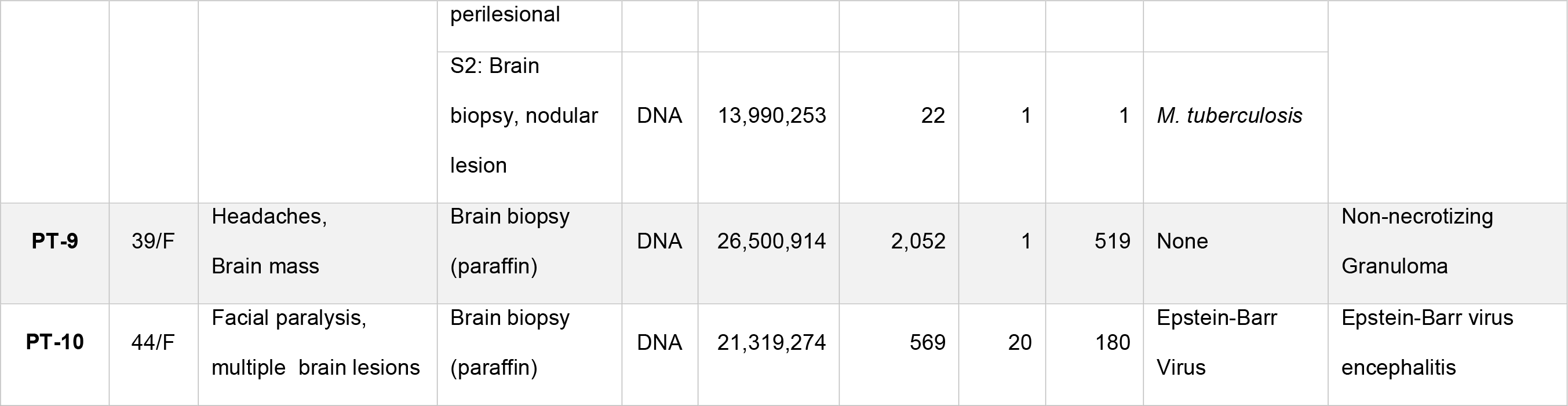
Summary of findings in ten patients from microbiome next-generation sequencing. Patients are numbered consecutively based on time of evaluation and diagnosis, with PT-1 being the first and PT-10 the last. Show are total numbers of NGS reads and numbers of bacterial, viral, and fungal reads. Sequencing-based diagnosis shows the microorganism indicated by sequencing as a possible cause of infection. Detailed read counts for all bacterial and viral species, as well as the number of human reads per sample, are found in Supplementary Tables 1-3.

### Computational processing

All reads were run through the Kraken system^12^ which compared them to a database containing the human genome (version GRCh38.p2), 2,817 bacterial genomes (representing 891 distinct species), 4,383 viral genomes (2,963 species), and 14 genomes of eukaryotic pathogens. The total size of the Kraken database in this study was 97 gigabytes. Each Kraken report was analyzed separately, and reads matching potential causative agents (bacteria or viruses) were extracted from the sequence file and re-aligned using the more sensitive BLASTN^13^ aligner against the comprehensive NCBI nucleotide database (nt), which contains many thousands of draft genomes and partial sequences in addition to the finished genomes in the custom Kraken database used here. Only reads whose BLAST matches hit the same species were considered further. Reads that hit different species were considered uninformative and were excluded from further analysis. For all of the positive diagnoses reported here, the BLAST results confirmed the species identified by Kraken, although the best-matching strain was sometimes a different (draft) genome (See Supplemental Methods for additional details).

## Results

### Sequencing data

The sequenced biopsies generated between 8.3 and 29.1 million reads per sample, summarized in Table 1. The vast majority of reads in all cases were human, as expected. (Note that because the human genome is approximately 1,000 times larger than most bacterial genomes, a mixture with equal numbers of human and bacterial cells will yield 99.9% human DNA.) After filtering to remove common contaminants and vector sequences, we then ranked the microbial species and considered only those that represented more than 10% of the reads and also ranked among the top three species as possible infectious agents (Figure 1). The number of bacterial and viral reads varied from fewer than 20 to 25,000. NGS successfully identified with high confidence the infectious agent in three out of 10 patients and possible pathogens in two patients. In the remaining five patients with negative or non-specific findings, NGS studies further helped to rule out presence of infectious pathogens in the disease process and supported therapeutic decisions in 3 patients. In two cases, diagnoses of non-infectious processes (e.g., glioblastoma and rapidly progressing astrocytoma) were identified by histopathological techniques.

Cases with a high degree of diagnostic confidence and positive pathogen identification (**See supplementary information for detailed clinical summaries**

**Figure 1.**
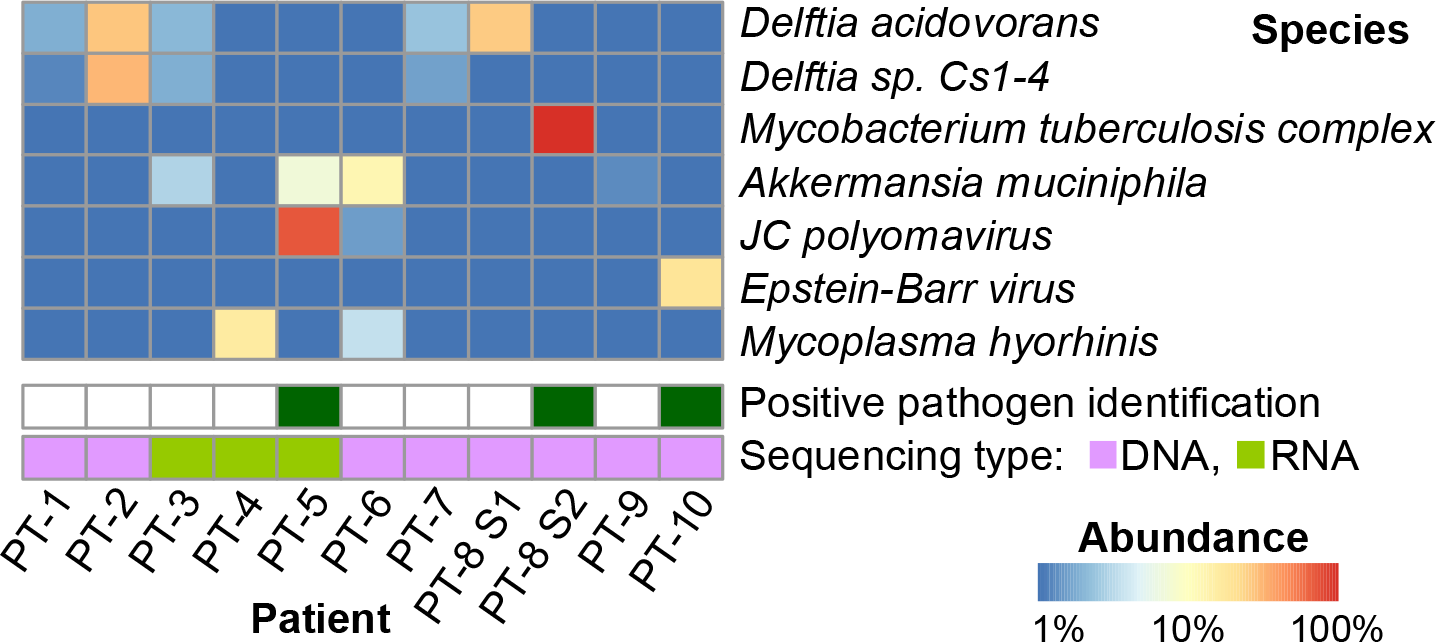
Heatmap showing the top microbial species in each of the eleven samples. All microbial species that explain 10% or more of microbial reads (excluding contaminants) in any sample are shown. Shaded squares indicate the relative proportion of reads from each species in that sample, ranging from blue (1%) to red (100%). Green boxes across the bottom indicate the three samples for which the sequence-based diagnosis was independently confirmed.

### Patient PT-8 (Pathogen identified: *Mycobacterium tuberculosis*)

A 67-year-old woman with osteomyelitis and multiple nodular lung lesions presented with multifocal brain and spinal lesions. Brain MRI showed progression of multiple nodular enhancing lesions throughout the brain compared to prior imaging (Figure 2). Biopsies from the perilesional (S1) and a nodular brain lesion (S2) were obtained for pathology, microbiology, and NGS studies. Two DNA sequencing runs yielded 15M reads from sample S1 and 14M reads from sample S2. These runs yielded the fewest microbial reads of any of the patients in our study: 18 and 22 bacterial reads, and only one to six viral and fungal reads, respectively, for samples S1 and S2. Nonetheless, a clear finding emerged for sample S2: 15 reads from *Mycobacterium tuberculosis.* Despite the small absolute number of reads, this species explained 68% of the bacterial reads detected. We manually confirmed the sequence assignments using Blast^13^ to align them against the NCBI nt database. We then re-aligned all reads against one specific genome, *M. tuberculosis* 7199-99 (accession NC_020089.1) using Bowtie2 with sensitive local alignment settings^14^. This procedure yielded 34 reads that were randomly distributed along the *M.* tuberculosis genome. Histopathological studies of the corresponding S2 sample showed necrotizing granulomas although studies with AFB, GMS and other stains failed to identify any microorganism (Figure 2). The patient responded rapidly to anti-tuberculous treatment and was discharged.

**Figure 2.**
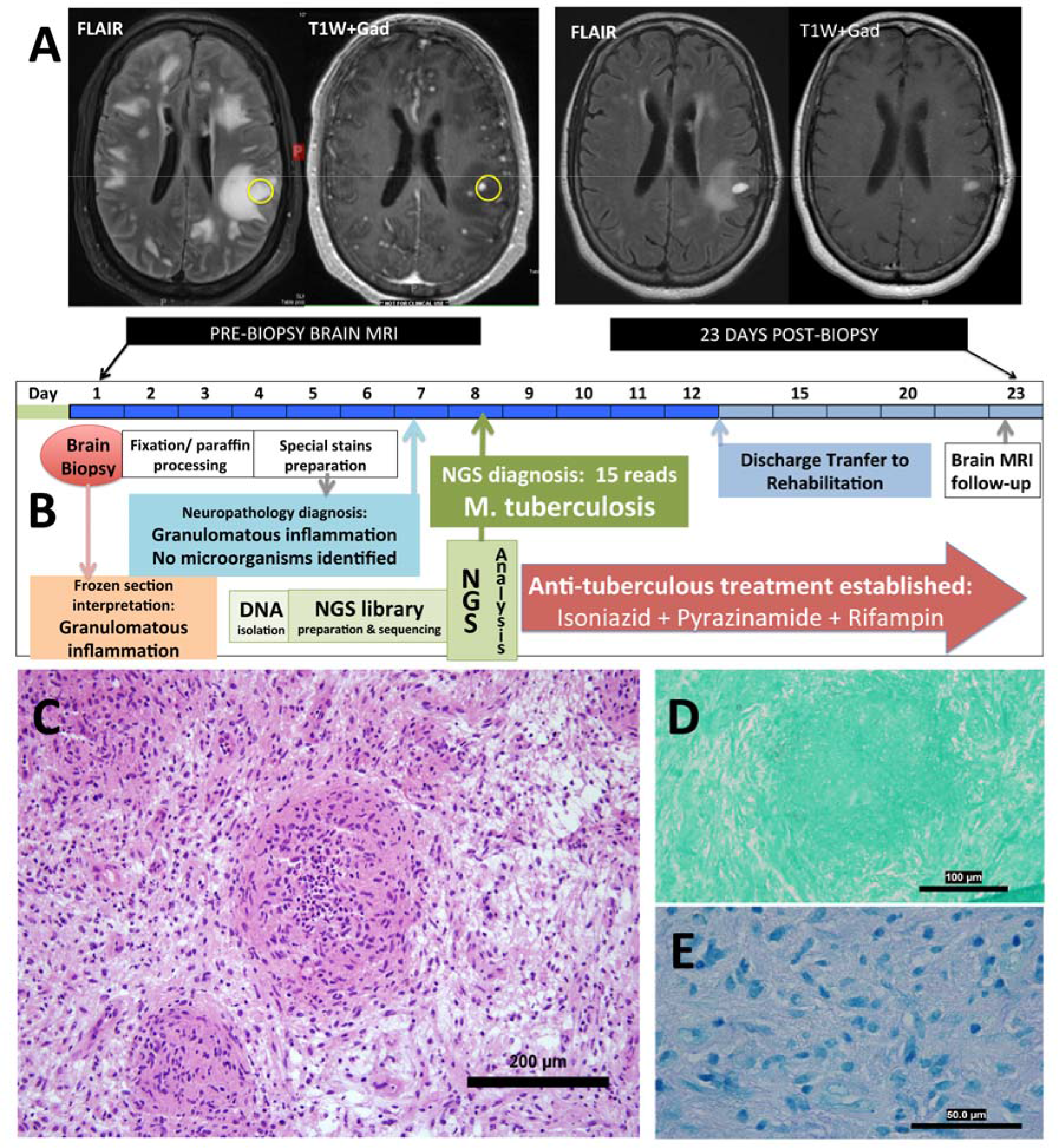
A patient with multifocal nodular lesions diagnosed with CNS tuberculosis. A) Pre-biopsy brain MR imaging in patient PD-8 showed multifocal subcortical and deep white matter lesions in FLAIR MR images. Several focal nodular enhancing lesions were seen in TlW+Gadolinium MR images. A brain biopsy targeted a nodular lesion in the left posterior fro ntalp arie tal region (yellow circles). Subsequent brain MR images demonstrated a dramatic improvement of the lesions 23 days after biopsy and 15 days after instauration of the anti-TB medications. B) Diagrammatic representation of the time-profile for biopsy processing for neuropathology and NGS diagnosis in PD-8. Although histological studies were able to establish the presence of a granulomatous inflammation immediately after biopsy, a more comprehensive neurohistological analysis completed 6 day after biopsy failed in demonstrating infectious pathogens. NGS diagnosis was established *7* days after biopsy confirming the presence of *M*. tuberculosis, a finding that was used to support the treatment with anti-tuberculous medication and ruled out the presence of Nocardia infection. C) Histological demonstration of granulomatous inflammation in H&E stains. D) GMS stain was negative for fungal species. Similarly, ZN stain for AFB (E) was negative.

### Patient PT-5 (Pathogen identified: JC Polyomavirus)

A 52-year-old man had been admitted for evaluation of right lower extremity weakness and gait disturbance, which evolved to right hemiparesis and a simple partial motor seizure. A brain MRI showed focal atrophy of the left post-central and adjacent superior frontal gyri and a focal white matter lesion (Figure 3A). PCR CSF studies for viruses and bacterial cultures were negative although a pre-biopsy CSF PCR was reported inconclusive or with “low DNA levels” for JCV. A biopsy of the white matter lesion was performed. RNA sequencing of the biopsy yielded 26.9 million reads, of which 25.9 million were human (Supplementary Table 1). Analysis found a very strong presence of JC Polyomavirus (JCV) with 8,944 out of 8,954 reads from all viruses. Although many bacterial species were detected, JCV was the most abundant species in terms of the number of reads, despite its small genome size. The whole genome of JCV was covered by the reads, at an average depth over 200 (Figure 3B). We therefore concluded that the sequence data showed strong support for infection with JCV, a known cause of PML^15^. Pathology results showed marked astrogliosis and intra-nuclear inclusions in oligodendrocytes (Figure 3C) and positive immunostaining for SV40 T antigen (a surrogate for JCV), confirming the diagnosis (Figure 3D).

**Figure 3.**
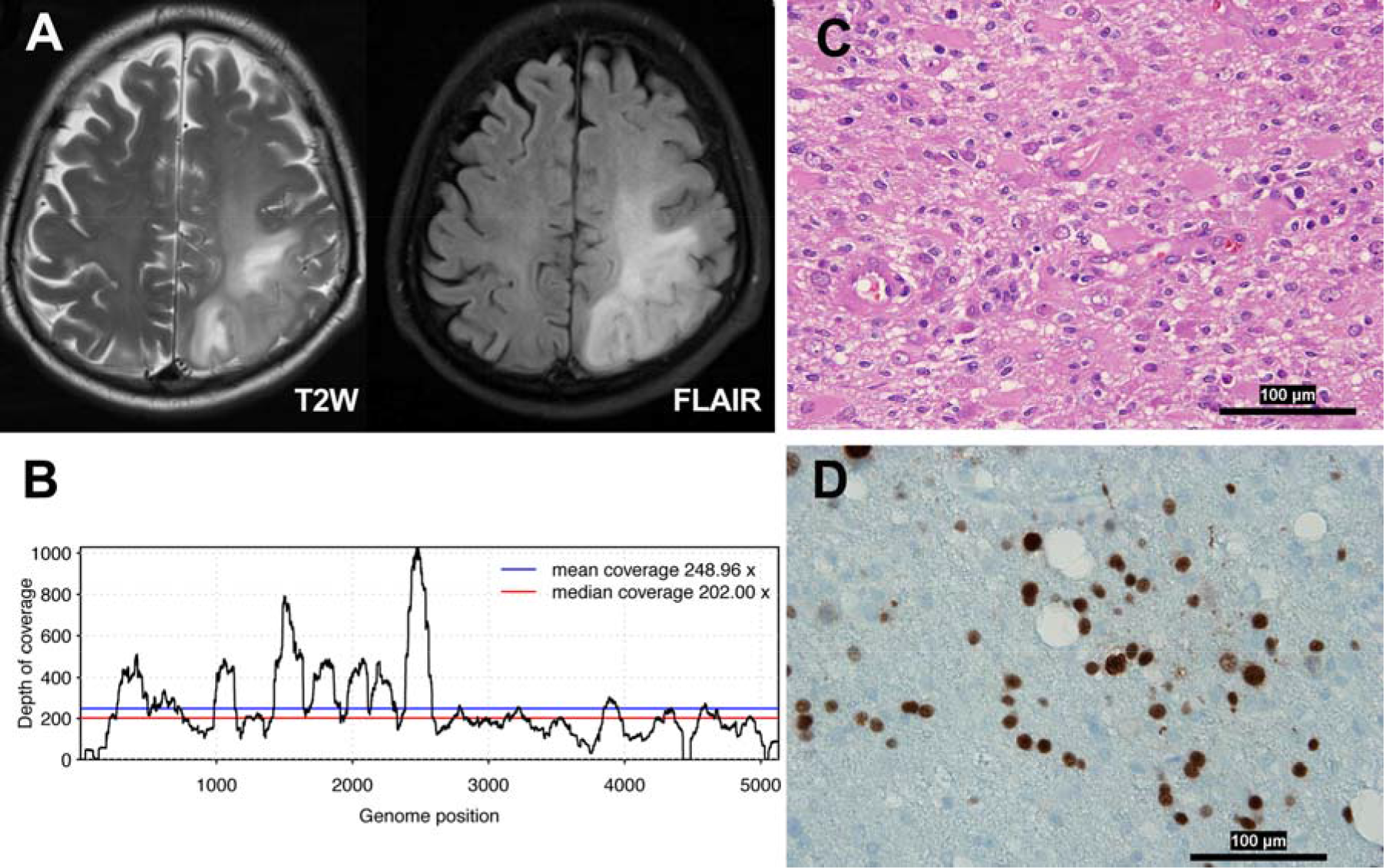
Brain imaging and neuropathological studies in patient PT-5 diagnosed with progressive multifocal leukoencephalopathy. A) MRI in patient PT-5 showed an extensive area of signal abnormalities in both T2W and FLAIR MRI sequences which involved the white matter of the left posterior region of the frontal-parietal lobes. B) NGS analysis revealed 8,944 JC polyomavirus reads out of 8,954 reads from all viruses. The whole genome of JC polyomavirus was covered by the reads, at an average depth over 200. The NGS findings were consistent with a diagnosis of PML. C) Histological changes observed in H&E stain which demonstrated extensive gliosis, presence of gemystocytic astrocytes and mild inflammatory reaction. D) Immunostaining with SV-40 antibodies, a surrogate for JC virus identification, showed several immunopositive nuclei as confirmation of JCV infection. These findings confirmed the diagnosis of PML.

### Patient PT-10 (Pathogen identified: Epstein-Barr virus)

A 44-year-old woman with a history of organ transplants and immunosuppression presented for evaluation of facial paralysis. Brain MRI showed at least three enhancing lesions in both hemispheres, one of them with appearance of a “ring enhancing” lesion which resembled CNS toxoplasmosis (Figure 4A-C). Assessment of CSF for known opportunistic infections was negative as well as CSF flow cytometry for malignancies. A biopsy of the left parietal lobe lesion showed granulomatous, lymphohistocytic inflammation and focal necrosis (Figure 4D). Paraffin sections were processed for NGS, which yielded 21.3 million reads (Table 1), of which 21 million were human and ∽216,000 were vector or synthetic controls. Only 569 reads were bacteria, all matching known skin bacteria or contaminants. Twenty reads matched viruses, of which 18 (90%) matched Epstein-Barr virus (EBV). A validation test using in situ hybridization for EBV-encoded RNA (EBER) confirmed EBV infection (Figure 4E). This case shows many similarities to a previous report^16^ of EBV-induced brain lesions in immunosuppressed patient following organ transplantation.

**Figure 4.**
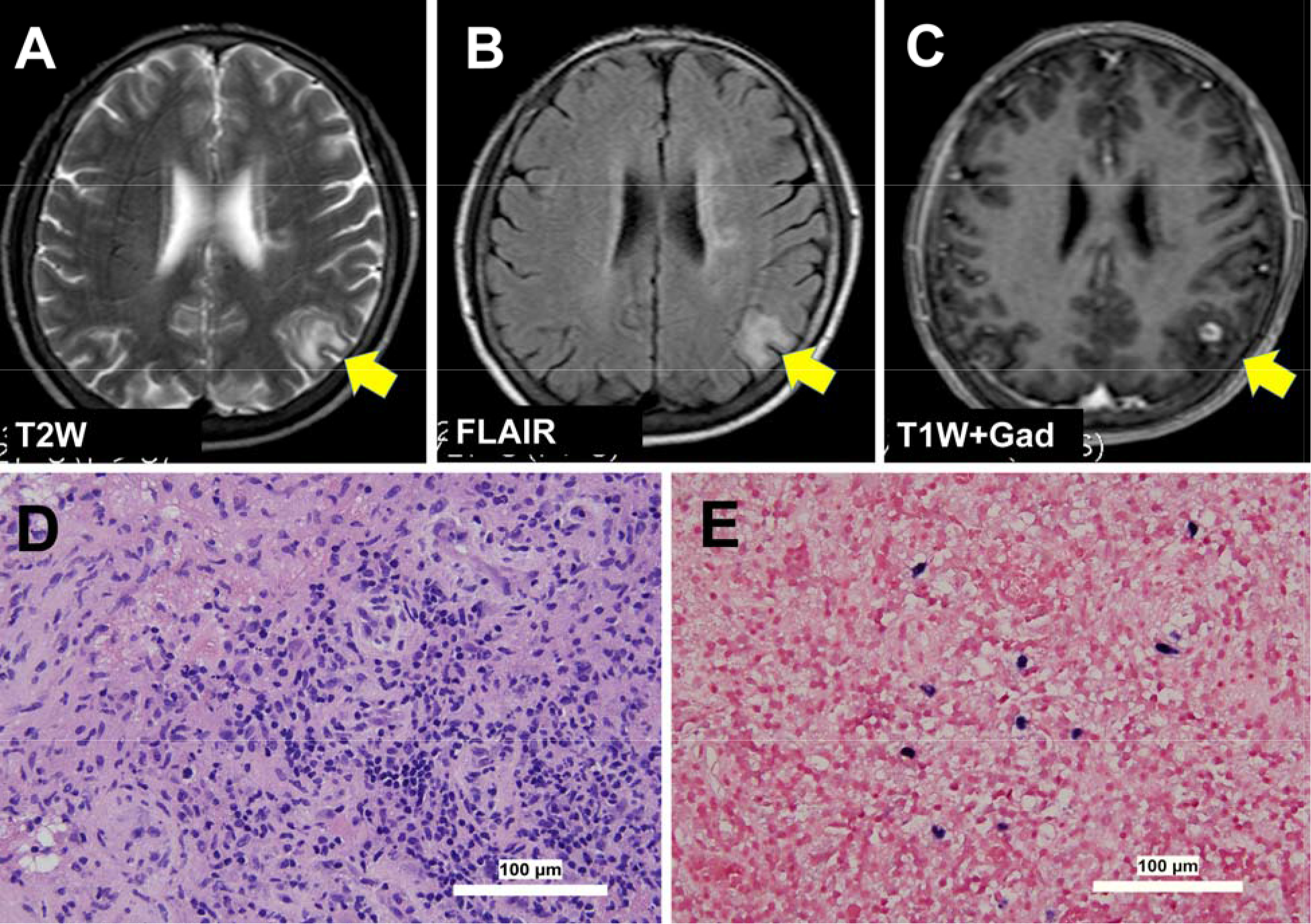
Brain imaging and neuropathological demonstration of EBV encephalitis in patient PT-10. A-C) Brain MRI showed a focal area of cortical and subcortical signal intensity abnormality. T1W sequences enhanced with gadolinium showed a “ring enhanced” lesion with perilesional edema which was the target for the brain biopsy D)Histological demonstration of a granulomatous and lymphohistocytic inflammation with foci of necrosis (H&E stain) in patient PD-10. E) In-situ hybridization for EBV encoded RNA (EBER) in the brain biopsy tissues which validated the presence of EBV as it had been established by NGS and confirmed the diagnosis of EBV encephalitis.

### Cases with candidate pathogen identification (not confirmed by a second method). Patient PT-2 with a Tolosa-Hunt-like syndrome with focal pachymeningitis suspected to be associated with chronic dural infection by Delftia

A 69-year-old man developed left-sided ptosis and retro-orbital headache after cataract and lens implantation surgeries. The initial symptoms were followed by decreased vision, horizontal diplopia, ophthalmoplegia and facial numbness. He was diagnosed with Tolosa-Hunt syndrome and treated with dexamethasone. A brain MRI demonstrated pachymeningeal enhancement localized in the medial aspect of the left middle cranial fossae, orbital apex, left cavernous sinus, Meckel’s cave, and foramen ovale (Figure 5A). Because of progressive brain changes on MRI and persistence of symptoms despite steroid treatment, the patient underwent biopsies of the skull base mass. Pathological studies of the dural lesion showed chronic inflammation, fibrosis (Figure 5C) and a moderately dense inflammatory infiltrate. Microbiological studies for infection were negative. NGS studies of the dura biopsy showed *Delftia acidovorans* (or possibly *Chryseobacterium taeanense*, which is very closely related to *Delftia*) and *Corynebacterium kroppenstedtii*. Although *Delftia* was seen in other samples, the relative proportion of *Delftia* reads in this patient was much higher (45% of non-human and non-contaminant reads, see Supplementary Table 2) than in any other patient. It was concluded that the pachymeningeal reaction was likely associated with a chronic evolving infection possibly triggered by the cataract removal and intra-ocular lens implantation. Treatment with a combination of IV ceftriaxone plus oral moxifloxacin produced clinical improvement and decreased dural enhancement in brain MRI (Figure 5B).

**Figure 5.**
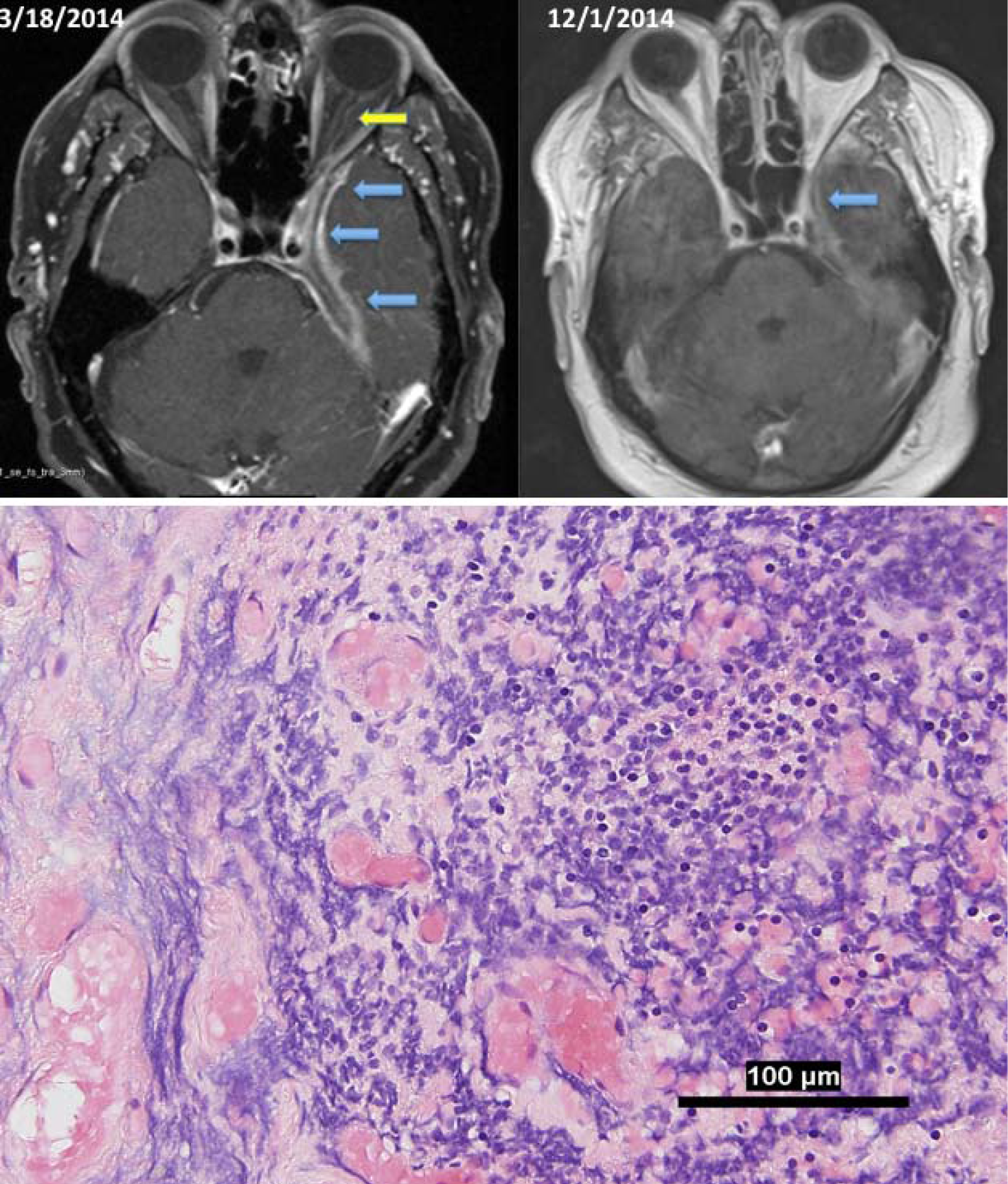
A patient with focal pachymeningitis and Tolosa-Hunt-like syndrome. A) Neuroimaging studies in patient PT-2 demonstrating the presence of pachymeningeal and leptomeningeal enhancement (blue arrows) localized in the medial aspect of the left middle cranial fossae extending to the orbital apex with involvement of the dural margin of the left cavernous sinus, Meckel’s cave, and foramen ovale Enhancement of the left optic nerve dura (yellow arrow) is also noted in the 3/18/2014 MRI. Neuroimaging studies following 5 weeks of antibiotic treatment (12/1/2014) showed decreased meningeal and dural enhancement along the anteromedial left temporal lobe margin and tentorial leaflet as well as optic nerve dural sheath. B) Histopathological studies of the dura and adjacent bone showed chronic inflammation comprised by lymphocytes, macrophages, and a few plasma cells (H&E stain).

### Patient PT-7: A patient with Fanconi’s anemia and neurodevelopmental disorder with new onset weakness and cerebritis suspected to be associated with *Lactococcus lactis* infection

This 19-year-old male had a previous history of dysgenesis of the corpus callosum and Fanconi’s anemia developed new-onset right-sided weakness and seizures. A brain MRI demonstrated a left-sided brain mass (Figure S1) and a brain biopsy suggested the diagnosis of lymphoma. He was treated initially with IV and intrathecal dexamethasone via an Ommaya reservoir. The patient deteriorated clinically and was transferred to our institution. A second biopsy of the brain lesion showed chronic inflammation (Figure S1) and acute coagulative necrosis. Studies for bacterial and fungal infection were negative. NGS studies of the biopsy did not reveal a clear candidate, but did show an unusually high presence of *Lactococcus lactis*, a common additive in dairy products that rarely causes human infections. 244 reads mapped to the genus *Lactococcus*, 201 of which were specific to *Lactococcus lactis cremoris*. NGS analysis also returned the second-highest number of viral reads (74) after PT-5 (Table 1), 58% of which matched to *L. lactis* phage species that we did not observe in other samples.

### Cases with non-specific or negative findings that were clinically useful

Sequencing yielded no specific findings to support a diagnosis of infection in 5 cases (Table 1); however the sequencing results did help to rule out concerns about an active infection in 3 cases. In case PT-3, a 23-year-old woman with status epilepticus following a febrile illness was diagnosed with *febrile infection related epilepsy syndrome* (FIRES),^17^, after extensive assessment and NGS studies were negative for viruses, and other pathogens. In other two cases PT-4 and PT-9, suspected to have granulomatous inflammation or sarcoidosis based on brain MRI and biopsy findings, NGS showed no evidence of specific infection, a finding that helped to support a decision for treatment with steroids. This therapy was associated with improvement in both cases. In two cases, neuropathological studies demonstrated that the disease process was associated with glial tumors despite initial concerns about CNS infections. In case PT-1, NGS studies showed several species of bacteria (Supplementary Table 3) all of which were suspected to be contaminants. In case PT-6, DNA sequencing identified 2,854 reads as bacterial and 17 as viral. While the bacterial reads did not reveal a potential pathogen, 15 of the 17 viral reads mapped to JCV, a finding considered to be incidental, although JCV has been implicated in the pathogenesis of astrocytomas in non-human primates^18^.

## Discussion

We describe the power of large-scale unbiased sequencing along with computational analysis as a diagnostic tool to establish the presence or absence of active infection in brain biopsies from patients with neuroinflammatory disorders suspected to be associated with pathogenic microorganisms. The current clinical standards for neuropathological diagnosis of infections, which include histological, immunological and selected molecular techniques (e.g., PCR, 16S rRNA, in-situ hybridization), have limited power as only a few infections of the CNS have specific histological hallmarks, and the diagnostic tools are limited even when specific neurological infectious disorders are suspected. We sought to test the effectiveness of metagenomic sequencing in a clinical setting for the analysis of brain biopsies from patients suspected to have CNS infections. In our study, NGS facilitated the identification of a pathogenic agent in three patients for whom the presence of the agent was further validated by histology, immunohistology, and/or molecular techniques. Potential pathogenic microorganisms were identified, though not confirmed, in two additional patients. In these five patients, standard diagnostic studies had failed to define etiological agents. In case PT-8, in which *Mycobacterium tuberculosis* was identified by NGS, the findings of sequencing studies helped to establish proper treatment that led to clinical improvement. Diagnostic confidence was strengthened by the histopathological findings of necrotizing granuloma and supported by the infrequency with which *M*. tuberculosis is a contaminant. Similarly in cases PT-5 and PT-10, the confidence in the findings of JCV and EBV was strengthened by immunocytochemical and *in-situ* hybridization techniques respectively. In those particular cases, NGS provided further support to determine the pathogenic role of these viruses. Although PCR studies for EBV and JCV in CSF are well established in clinical practice for evaluation of patients who have risk factors for opportunistic infection, there are specific clinical situations in which diagnostic tests such as brain biopsies are required. For example, the presence of EBV DNA in CSF of immunocompromised patients should be interpreted with caution as EBV may co-exist with other opportunistic infections^19^. In case PT-10 for which CSF PCR studies were negative, the finding of EBV by NGS and further validation by EBVAR *in-situ* hybridization helped to establish a definitive diagnosis of EBV-associated encephalitis, not only by demonstrating the presence of EBV in the inflamed tissue but also by excluding opportunistic infections such as toxoplasma which had been suggested by brain MRI. Similarly, the value of metagenomic sequencing in the diagnosis of PT-5 was demonstrated by the inconclusiveness of two pre-biopsy qualitative JCV-PCR studies in CSF,. Although immunocytochemistry and other molecular techniques (e.g., PCR) might have helped to establish the diagnosis of PML in this patient, case PT-5 represents a good proof-of-concept that NGS studies are useful and may provide better characterization of the JCV genetic variants useful for prognosis^20-21^.

Although a high degree of certainty could not be achieved in some cases, NGS provided information that facilitated a better understanding of the high-or low likelihood of CNS infection in the other 5 patients. In two cases, suspected microorganisms were identified although their pathogenic role could not be confirmed by histology, cultures or direct pathogen visualization. However, the high degree of suspicion for infection as the cause of the neuroinflammatory disorders prompted antimicrobial therapy. In case PT-2, reads from *Delftia acidovorans* and *Corynebacterium kroppenstedtii*, normal commensals of skin raised concern that these organisms were playing a pathogenic role in this patient’s pachymeningitis, particularly after progressive neurological problems despite empiric steroid treatment. Clinical improvement and reduction in the pachymeningeal inflammation as demonstrated by MRI evaluation suggest a chronic infection likely facilitated by the trafficking of commensal bacterial species into the duramater via ocular structures following the surgical procedure. Interestingly, in patient PT-7, who had a history of immunosuppression and a left hemisphere lesion suspected to be a lymphoma, NGS studies found not only a suspected pathogenic agent (*L. lactis cremoris*), but also a high number of reads from *L. lactis* phages, which were not observed in other cases. Because validation of *L. lactis cremoris* as a pathogenic organism could not be verified by other methods, the NGS findings were only suggestive of the potential pathogenic role of this bacteria. Although rare, *L. lactis cremoris* has been implicated in brain abscesses, brain empyemas and other infections in the brain^22-24^, we interpreted that this patient may have had chronic cerebritis associated with *L. lactis cremoris*, and his improvement after antibiotic treatment indirectly supports this finding.

Whereas microbiome analysis of brain biopsies was able to identify definite or possible pathogenic microorganisms in five of 10 cases, no pathogenic species were identified in the other five cases. However, NGS studies in three of those cases supported the conclusion that a brain infection was not present, findings that were also very valuable for understanding the neuropathological processes that affected such patients. In case PT-3, a young patient with status epilepticus after a febrile illness, the failure to demonstrate a pathogen helped to determine her neurological disorder as most likely associated with FIRES, a rare condition of uncertain etiology in which infection is suspected to be the triggering factor^17^. In another two cases, PT-4 and PT-9, NGS showed no clear evidence of pathogenic microorganisms associated with the granulomatous inflammation noted on brain biopsies. We therefore concluded there was no active infection, a conclusion that was further supported by the beneficial effect of steroid treatment. These two cases demonstrate the way in which negative NGS findings may help to determine treatment approaches, as the presence of granulomatous inflammation always raises concerns about the presence of slow-growing or atypical bacteria or fungal infections in which the use of steroids would be deleterious. Clearly, microbiome analysis of brain tissues has limitations, and the absence of pathogenic microorganisms should be interpreted with caution. It is possible that the timing of the brain biopsy may not coincide with the period of active infection, and thus inflammatory changes could represent reactive immune responses to an earlier infection. Negative NGS studies in patients with neuroinflammatory diseases may also suggest the cause of such disorders are due to post-infectious or autoimmune inflammatory responses rather than a direct result of active infection.

Although microbiome studies of brain biopsies shows considerable potential for detecting pathogenic and non-pathogenic microorganissms, contamination is always a concern as brain biopsies are subject to exposure to contaminants derived from the human body, the operating room, and tools used during biopsy procedures as well as carrier containers or the laboratory environment. In general, however, we expect a lower burden of contamination from environmental and other organ sources because the brain is isolated by the blood-brain barrier or meninges serving as a physiological seal protecting the CNS from the microbiome of other organs. Sequence-based results for patients in which NGS studies were negative or inconclusive, as well as in cases supporting specific diagnoses, contained common skin bacteria and other contaminants that were not considered clinically relevant (Supplementary Table 2). Most species were recognizable as common skin bacteria or possible contaminants from other sources. While we cannot rule out these species as pathogens, their presence in multiple samples collected from different patients widely spaced apart in time indicates that they are not likely to be causally related to the patients’ symptoms. For example, we observed the skin bacterium *Propionibacterium acnes* in almost all samples, illustrating the difficulty of capturing a pure sample without any microbial spillover from the intervening tissues. These findings illustrate the difficulty and complexity of identifying pathogens in human samples, which unavoidably contain some bacterial species because the human body is not a sterile environment. Virtually all sampling methods including brain biopsies must penetrate the skin, which is colonized with multiple bacteria including in sub-epidermal layers^25^, meaning that superficial cleansing will not remove them. In addition, surgical procedures which include the use of irrigation solutions as well as processing methods (e.g., DNA extraction kits and other sequencing reagents) often contain bacterial DNA that may be captured by NGS methods especially when the concentration of microbial DNA is low^26^.

An important limitation of computational analysis in microbiome studies is the size and completeness of genome sequence databases. Although thousands of bacterial and viral species have been sequenced, many others including human pathogens may yet be discovered, and unknown pathogens cannot detected until their genomes are sequenced. Eukaryotic pathogens represent a larger challenge: many of these remain to be sequenced, and even those that have been sequenced are available only in draft form, with many gaps and errors in the publicly available genomes. The addition of new and more complete genomes to public archives, which continue to grow rapidly, will steadily increase the power of computational analysis to detect pathogens from sequence data.

In summary, we described here the power of large-scale unbiased sequencing along with computational-based metagenomic analysis in the study of brain biopsies. Our study opens the possibility for more extensive use of NGS techniques in the identification of pathogens in patients with suspected CNS infections. Microbiome analysis of CNS tissues may therefore serve as a powerful tool in the identification of infective agents triggering neuroinflammatory disorders.

## Acknowledgements

We thank Drs. Arun Venkatasen, Matthew Ippolito and Robert McMillan for contributing ideas and for patient care. This research was supported in part by NIH under grant R01 HG006677 (S.L.S.), by the U. S. Army Research Office under grant number W911NF-14-1-0490 (S.L.S.) and by The Bart McLean Fund for Neuroimmunology Research-Johns Hopkins Project Restore (CAP).

## Supplementary Information

Supplemental Material includes supplementary clinical summaries, additional methods and data, and the figures and tables whose captions are given below.

**Supplementary Figure S1.** A patient with history of Fanconi’s anemia and brain malformation with a left hemispheric mass.

**Supplementary Table S1.** Percentages and raw counts of the number of reads found in each of the 11 samples, for all species that comprised at least 1% of any of the samples. Species with fewer than 5 reads in all samples are reported in aggregate as “Other microbial reads.” Values shown here were computed after removing human reads, known vector and adapter sequences, and the three most common contaminants: E. coli, S. cerevisiae, and P. acnes. Note that samples PT-8.S1 and PT-8.S2 came from two sites on the same patient.

**Supplementary Table S2.** Percentages and raw counts of the number of reads found in each of the 11 samples, for all species that comprised at least 1% of any of the samples. Values shown here were computed after removing human reads, known vector and adapter sequences. Species with fewer than 5 reads in all samples are reported in aggregate as “Other microbial reads.” As shown in the first three rows, the species E. coli, S. cerevisiae, and P. acnes dominate many of the samples, and were removed to produce the values shown in Supplementary Table 1.

**Supplementary Table S3.** Read lengths and total numbers of human and artificial reads for each sample.

**Supplementary Table S4.** Raw counts of the number of viral reads found in each sample.

**Supplementary Data.** Taxonomic summary of all Kraken classifications for patient PT5.

## References

1. Glaser CA, Honarmand S, Anderson LJ, et al. Beyond viruses: clinical profiles and etiologies associated with encephalitis. Clinical infectious diseases: an official publication of the Infectious Diseases Society of America 2006; 43: 1565–1577.

2. Barzón L, Lavezzo E, Militello V, Toppo S, Palu G. Applications of next-generation sequencing technologies to diagnostic virology. International journal of molecular sciences 2011; 12: 7861–7884.

3. Yin L, Liu L, Sun Y, et al. High-resolution deep sequencing reveals biodiversity, population structure, and persistence of HIV-1 quasispecies within host ecosystems. Retrovirology 2012; 9: 108.

4. Lecuit M, Eloit M. The human viróme: new tools and concepts. Trends in microbiology 2013; 21: 510–515.

5. Wylie KM, Weinstock GM, Storch GA. Viróme genomics: a tool for defining the human viróme. Current opinion in microbiology 2013; 16: 479–484.

6. Chan BK, Wilson T, Fischer KF, Kriesel JD. Deep sequencing to identify the causes of viral encephalitis. PloS one 2014; 9: e93993.

7. Wilson MR, Naccache SN, Samayoa E, et al. Actionable diagnosis of neuroleptospirosis by next-generation sequencing. The New England journal of medicine 2014; 370: 2408–2417.

8. Naccache SN, Peggs KS, Mattes FM, et al. Diagnosis of neuroinvasive astrovirus infection in an immunocompromised adult with encephalitis by unbiased next-generation sequencing. Clinical infectious diseases: an official publication of the Infectious Diseases Society of America 2015; 60: 919–923.

9. Brown JR, Morfopoulou S, Hubb J, et al. Astrovirus VA1/HMO-C: an increasingly recognized neurotropic pathogen in immunocompromised patients. Clinical infectious diseases: an official publication of the Infectious Diseases Society of America 2015; 60: 881–888.

10. Fremond ML, Perot P, Muth E, et al. Next-Generation Sequencing for Diagnosis and Tailored Therapy: A Case Report of Astrovirus-Associated Progressive Encephalitis. J Pediatric Infect Dis Soc 2015; 4: e53–57.

11. Hoffmann B, Tappe D, Hoper D, et al. A Variegated Squirrel Bornavirus Associated with Fatal Human Encephalitis. The New England journal of medicine 2015; 373: 154–162.

12. Wood DE, Salzberg SL. Kraken: ultrafast metagenomic sequence classification using exact alignments. Genome biology 2014; 15: R46.

13. Altschul SF, Madden TL, Schaffer AA, et al. Gapped BLAST and PSI-BLAST: a new generation of protein database search programs. Nucleic Acids Res 1997; 25: 3389–3402.

14. Langmead B, Salzberg SL. Fast gapped-read alignment with Bowtie 2. Nature methods 2012; 9: 357–359.

15. Ferenczy MW, Marshall LJ, Nelson CD, et al. Molecular biology, epidemiology, and pathogenesis of progressive multifocal leukoencephalopathy, the JC virus-induced demyelinating disease of the human brain. Clinical microbiology reviews 2012; 25: 471–506.

16. Khalil M, Enzinger C, Wallner-Blazek M, et al. Epstein-Barr virus encephalitis presenting with a tumor-like lesion in an immunosuppressed transplant recipient. Journal of neurovirology 2008; 14: 574–578.

17. Pardo CA, Nabbout R, Galanopoulou AS. Mechanisms of epileptogenesis in pediatric epileptic syndromes: Rasmussen encephalitis, infantile spasms, and febrile infection related epilepsy syndrome (FIRES). Neurotherapeutics: the journal of the American Society for Experimental NeuroTherapeutics 2014; 11: 297–310.

18. Maginnis MS, Atwood WJ. JC virus: an oncogenic virus in animals and humans? Semin Cancer Biol 2009; 19: 261–269.

19. Martelius T, Lappalainen M, Palomaki M, Anttila VJ. Clinical characteristics of patients with Epstein Barr virus in cerebrospinal fluid. BMC infectious diseases 2011; 11: 281.

20. Ryschkewitsch CF, Jensen PN, Major EO. Multiplex qPCR assay for ultra sensitive detection of JCV DNA with simultaneous identification of genotypes that discriminates non-virulent from virulent variants. Journal of clinical virology: the official publication of the Pan American Society for Clinical Virology 2013; 57: 243–248.

21. Marshall LJ, Ferenczy MW, Daley EL, Jensen PN, Ryschkewitsch CF, Major EO. Lymphocyte gene expression and JC virus noncoding control region sequences are linked with the risk of progressive multifocal leukoencephalopathy. Journal of virology 2014; 88: 5177–5183.

22. Feierabend D, Reichart R, Romeike B, Kalff R, Walter J. Cerebral abscess due to Lactococcus lactis cremoris in a child after sinusitis. Clinical neurology and neurosurgery 2013; 115: 614–616.

23. Inoue M, Saito A, Kon H, et al. Subdural empyema due to Lactococcus lactis cremoris: case report. Neurologia medico-chirurgica 2014; 54: 341–347.

24. Topcu Y, Akinci G, Bayram E, Hiz S, Turkmen M. Brain abscess caused by Lactococcus lactis cremoris in a child. Eur J Pediatr 2011; 170: 1603–1605.

25. Grice EA, Segre JA. The skin microbiome. Nature reviews Microbiology 2011; 9: 244–253.

26. Salter SJ, Cox MJ, Turek EM, et al. Reagent and laboratory contamination can critically impact sequence-based microbiome analyses. BMC biology 2014; 12: 87.

